# Protein semi-synthesis enables real-time optical tracking of intracellular conformational changes during sodium channel inactivation

**DOI:** 10.64898/2026.08.24.746605

**Authors:** Laurie Peverini, Michelle Nilsson, Iacopo Galleano, Vita Sereika-Bejder, Emma Krindel Beyer, Lukas Morten Fagerlund, Janne Møgelhøj Colding, Gerbrand van der Heden van Noort, Kristian Strømgaard, Stephan Alexander Pless

## Abstract

Dynamic conformational changes in intracellular domains fundamentally affect the function and pharmacology of many membrane proteins. For example, sodium influx through the cardiac voltage-gated sodium channel (Na_V_1.5) is rapidly terminated through conformational changes that result in pore closure, a transition known as inactivation. Inactivation involves Na_V_1.5 intracellular regions, particularly the DIII-DIV linker containing the IFM particle (Isoleucine-Phenylalanine-Methionine) and its dysfunction is a major cause of cardiac arrythmias. However, the conformational changes involved in inactivation and their modulation by auxiliary proteins and clinically used drugs remain incompletely characterized, partly because live-cell, site-specific labeling of intracellular regions with small fluorescent dyes remains challenging. Here, we combine live-cell protein semi-synthesis with voltage-clamp fluorometry (VCF), to track intracellular conformational dynamics of the cardiac sodium channel Na_V_1.5 and monitor their voltage dependence and kinetics in real time. We identify intracellular conformational changes involved in both fast and steady-state inactivation of Na_V_1.5 and show that both lidocaine and auxiliary proteins affect the kinetics of conformational changes of the DIII-DIV linker. Our work establishes the combination of protein semi-synthesis and voltage-clamp fluorometry as a powerful approach to dissect intracellular conformational changes in membrane proteins.

## Introduction

Ion channels are transmembrane proteins capable of converting chemical, physical or mechanical stimuli into electrical signals through conformational changes that often propagate throughout their entire structure. However, the structure of their intracellular domains (ICDs) often remains unresolved and their conformational transitions are difficult to track with high temporal resolution in cellular environments. The contributions of ion channel ICDs to function and pharmacology of these important drug targets thus remain largely enigmatic. This is also true for voltage-gated sodium channels (Na_V_s), which respond to membrane depolarization with their voltage-sensing domains (VSD) undergoing an upward motion that drives a cascade of conformational changes. These result in opening of the central ion-conducting pore and thus the entry of sodium. Immediately after channel opening, Na_V_s undergo a conformational transition known as fast inactivation, which terminates sodium influx^1^. By contrast, steady-state inactivation (SSI) governs channel availability during prolonged depolarizations and can occur from closed states. In the case of Na_V_1.5 (hNa_V_1.5), which is predominantly expressed in cardiomyocytes, both forms of inactivation maintain normal cardiac excitability and their disruption through mutations or other modifications can cause severe arrhythmias^2^. Like other Na_V_s, hNa_V_1.5 is a monomeric pseudo-tetramer of around two thousand residues that assembles into four domains (DI-DIV), with six transmembrane helices each (S1-S6), connected by intracellular domains of various sizes and the N and C-terminal parts are intracellular^2^. Inactivation involves conformational changes in both transmembrane and intracellular domains^2^. This causes the IFM particle (Isoleucine-Phenylalanine-Methionine) in the DIII-DIV linker to dock into a hydrophobic pocket between DIII S5, DIV S5/S6 helices and DIV S4-S5 linker, resulting in allosteric pore closure^3,4^.

However, the precise nature of the intracellular conformational changes underlying hNa_V_1.5 inactivation, as well as their modulation by drugs or auxiliary proteins remain incompletely characterized due to challenges in tracking them in real-time in living cells. Additionally, and although numerous structures have been published^3,5–13^, the full gating cycle of Na_V_1.5 in a membrane environment has not been structurally resolved. The available cryo-EM structures, determined in the absence of an electric field, are often difficult to assign to a defined conformational state. Thus, complementary methods such as voltage-clamp fluorometry (VCF)^14– 16^, which allows simultaneous live-cell tracking of conformational changes and functional state of the channel are of particular interest in this context^16^. To perform said VCF experiments it is necessary to fluorescently label the ion channel of interest, but this can be challenging when targeting intracellular domains of membrane proteins as they cannot be labelled with conventional cysteine-reactive fluorescent probes. Although engineered aminoacyl tRNA synthetase/tRNA pairs can be used for non-sense suppression-based methods to insert non-canonical fluorescent amino acids into ICDs, this only allows incorporation of moderately bright fluorescent probes with typically limited photostability and can lead to non-homogenous population of ion channels^17^. In order to develop a versatile methodology to site-specifically introduce a bright and photostable fluorescent probe into a membrane protein, we adapt our tandem Protein Trans Splicing (tPTS) approach to achieve site-specific fluorophore insertion^18,19^. This enables us to perform voltage-clamp fluorometry (VCF) recordings with a fluorescent probe incorporated into the hNa_V_1.5 DIII-DIV intracellular linker and to directly investigate its conformational changes in response to depolarizations and binding of drugs or auxiliary proteins. We show that that binding of auxiliary proteins and pore block by lidocaine, which both occur in distal protein domains, allosterically affect the kinetics of DIII-DIV linker conformational changes and thereby establish a powerful method to track conformational changes in membrane protein ICDs.

## Results

To obtain fluorescent hNa_V_1.5, we recombinantly express the N- and C-terminal fragments of hNa_V_1.5 (amino acids (aa) 1 to 1471 (N) and 1503 to 2016 (C)) and inject a synthetic peptide X (aa 1472 to 1502) in *Xenopus laevis* oocytes (see extended Methods). The N, C and X fragments are spliced by two orthogonal split inteins, *Cfa*DnaE and *Ssp*DnaB^M86^ (Fig 1A)^19–21^. Residue K1492 in X is conjugated to a rhodamine 110, allowing hNa_V_1.5 reconstitution with a fluorescent label in the α-helix upstream of the IFM particle (Fig 1B). Reconstituted hNa_V_1.5 (NX^Rho^C), but not the NC control, shows robust expression and wild type-like voltage dependence of activation (V_50(Act)_ = - 39.4 +/- 0.3 mV (WT) *vs* -41.3 +/- 0.8 mV (NX^Rho^C)), although SSI is altered (V_50(SSI)_ = -69.7 +/- 0.5 mV (WT) *vs* -58.1 +/- 0.6 mV (NX^Rho^C)), late currents are increased (1.3 +/- 0.9 % for NX^Rho^C *vs* - 1.4 +/- 0.3 % for WT, Fig 2C) and recovery from inactivation is accelerated (5.8 +/- 0.5 ms for NX^Rho^C *vs* 14.1 +/- 1.4 ms for WT, Fig 1C).

**Figure 1.**
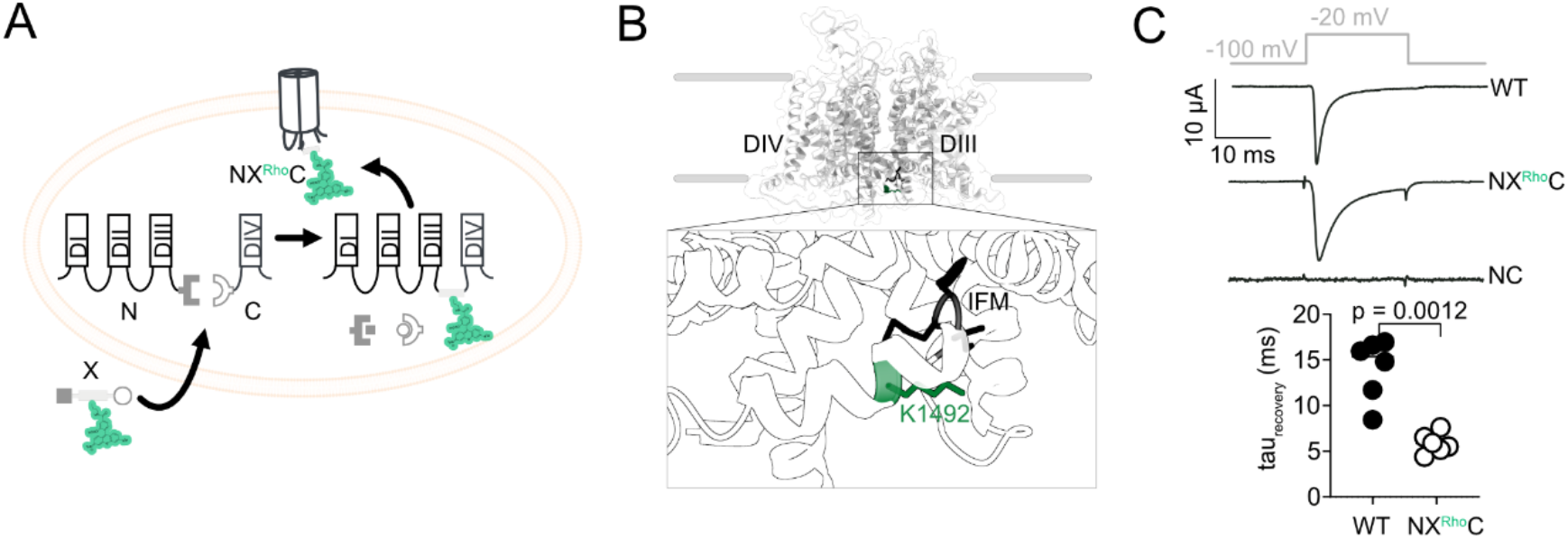
Development and expression of a fluorescent hNav1.5 (NX^Rho^C). **A**. Schematic of the tPTS-based approach to label the hNa_v_1.5 DIII-DIV linker. **B**. hNa_V_1.5 structure (adapted from PDB: 6lqa^7^) with inset showing the DIII-DIV linker with IFM motif (black) and K1492 labelling site (green). **C**. Currents for WT, NX^Rho^C and NC only (top) and time constants (tau) of current recovery (Welch’s t-test, bottom).

**Figure 2.**
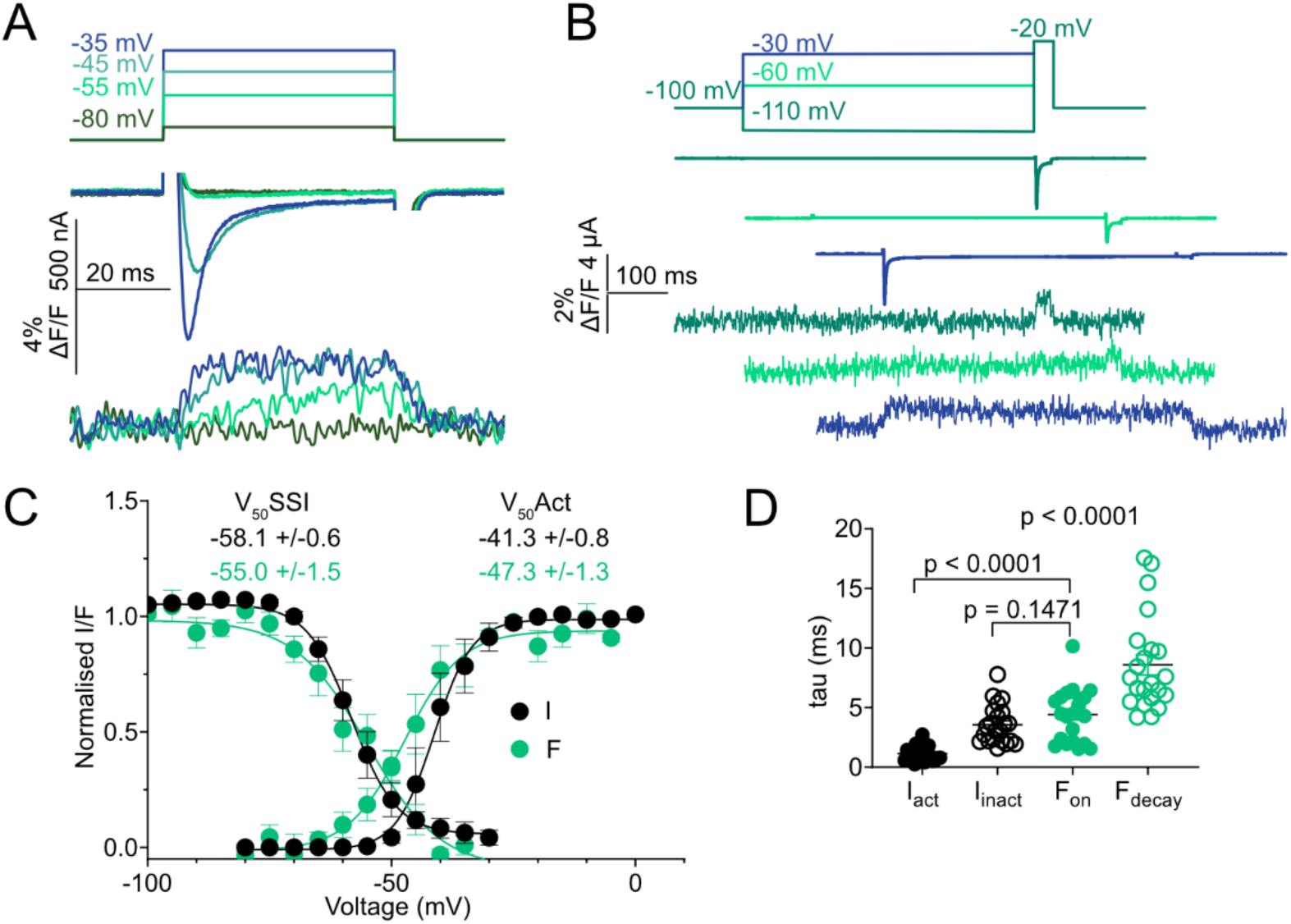
VCF recordings and characterization of NX^Rho^C. **A/B**. Color-coded voltage steps (top), current (I, middle) and fluorescence (F, bottom) recordings for activation (**A**) and inactivation protocols (**B**, traces are shifted horizontally for clarity). **C**. SSI (left) & activation (right) curves for I and F (mean +/- SEM, n=4-7). **D**. Tau values for I and F parameters obtained at -30 mV (Welch’s t-test, mono-exponential fits).

Strikingly, depolarization evokes fluorescence signal (ΔF) dequenching (Fig 2A), consistent with the fluorescent probe placed in the α-helix upstream of the IFM particle moving to a more hydrophobic environment as observed in atomic structures^3,22^. Importantly, ΔF is elicited at subthreshold voltages where small or no currents are recorded (Fig 2A), suggesting that ΔF is not triggered by channel activation, but potentially arise from transitions to pre-open or inactivated states. This is corroborated by the fact that V_50(Act)_ of ΔF is left-shifted by around 6 mV compared with V_50(Act)_ of current (Fig 2B/C). Furthermore, the onset of ΔF (F_on_) is slower than current activation (I_act_) (Fig 2D), and ΔF remains stable during the voltage step, while the current is rapidly inactivating (Fig 2A). By contrast, SSI protocols (Fig 2B) reveal that the voltage dependence is superimposed for current and fluorescence, and that F_on_ matches the current kinetics of fast inactivation (I_inact_) (Fig 2D). This suggests that the DIII-DIV linker movement tracks conformational changes related to hNa_V_1.5 inactivation.

Next, we investigate if protein-protein interactions (PPIs) at the C-terminal domain (CTD) or drug binding to the inner pore, which both affect inactivation, allosterically influence the DIII-DIV linker conformational changes. iFGF12A (intracellular fibroblast growth factor homologous factor 12A) is a cytosolic protein that binds to the hNa_V_1.5 CTD to reduce late currents and modulate SSI^23^. Co-expression of iFGF12A with NX^Rho^C reveals a similar fluorescence dequenching than without (Fig 3A) and it reduces the late current (Fig 3C), suggesting that the PPI remains partially preserved, although it does not alter SSI (Fig 3B), likely due to rhodamine labelling. SSI of current and ΔF are unaffected by iFGF12A co-expression (Fig 3B), but F_on_ kinetics are slower than those of I_inact_ and the decay of fluorescence (F_decay_) is slowed (Fig 3D/E).

**Figure 3.**
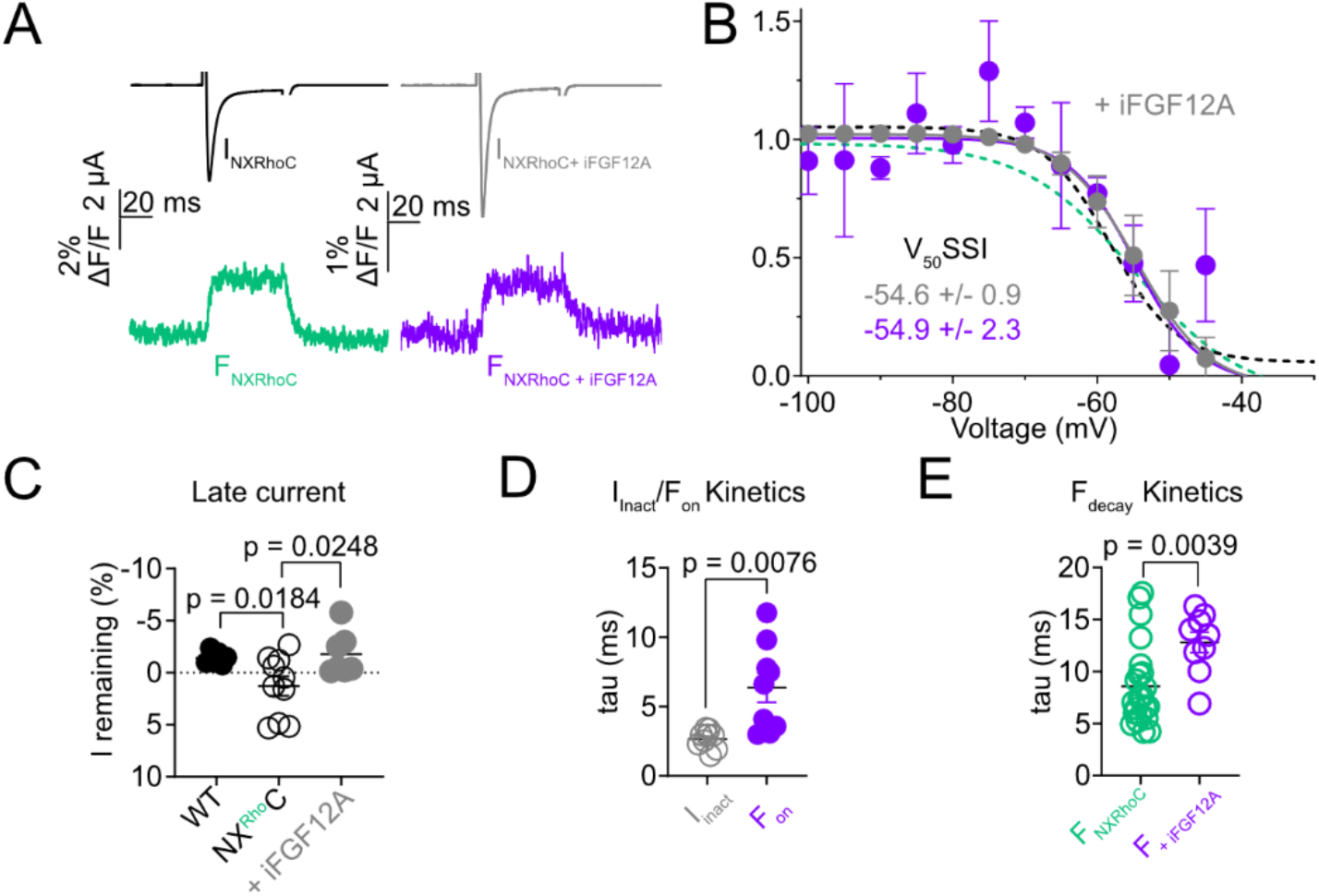
PPIs affect DIII-DIV linker conformational changes. **(A)/(B)** Current (I) and fluorescence (F) recordings (A) and SSI data (B) for NX^Rho^C +/- iFGF12A (+: grey: I, purple: fluorescence; -: black/green dashed lines; mean +/- SEM, n=3-6). **C**. Late current +/- iFGF12A. **D**. F_on_ versus I_inact_ kinetics at -30 mV. **E**. F_decay_ (F_off_) +/- iFGF12A at -30 mV. Welch’s t-test for **C, D** and **E**.

Finally, we expose NX^Rho^C to lidocaine, a class Ib anti-arrhythmic drug that blocks hNa_V_1.5 by binding preferentially to open and inactivated states^24,25^. We observe that depolarization causes a dequenching of the fluorescence also in the presence of lidocaine (Fig 4A). Application of lidocaine in a moderate use-dependent block regime (-30 mV at 3 Hz) reduces peak current more than ΔF amplitude (Fig 4B) and elicits no additional ΔF components (Fig 4A), suggesting that lidocaine does not *per se* interfere with DIII-DIV linker movements. Strikingly, F_on_ is faster than I_inact_, and F_decay_ is slowed by lidocaine in a concentration-dependent manner (Fig 4C/D). These findings highlight how PPIs and drug binding can allosterically modulate hNa_v_1.5 DIII-DIV linker movements.

**Figure 4.**
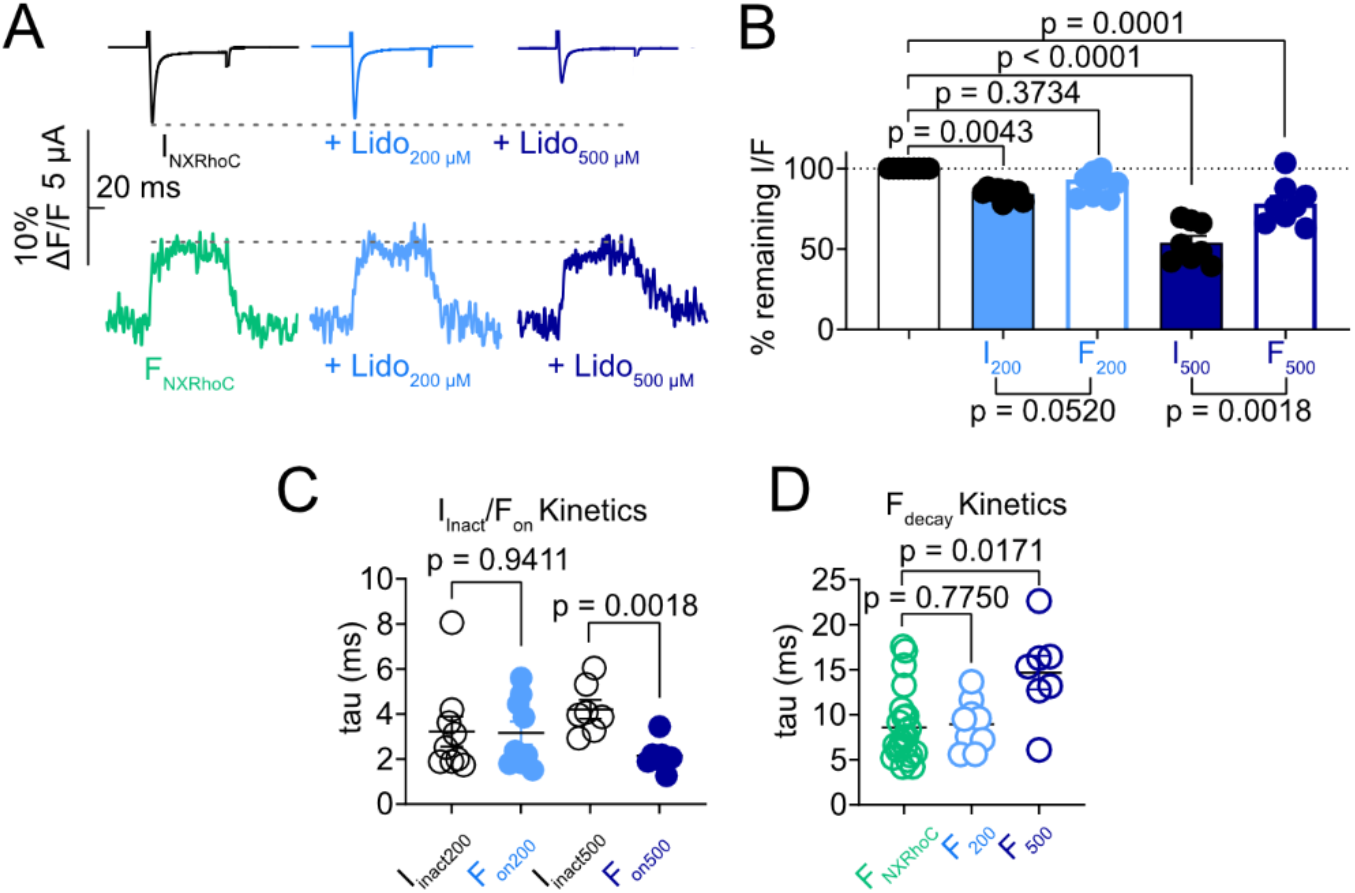
Lidocaine affects DIII-DIV linker conformational changes. **A**. Current (I) and fluorescence (F) recordings upon lidocaine application (200 μM light blue, 500 μM blue). **B**. Quantification of I (filled) and F (empty) remaining after 200/500 μM lidocaine application (one-way ANOVA with Dunnett’s test for maximal value, Welch’s t-test for I vs. F). **C**. F_on_ versus I_inact_ kinetics at -30 mV. **D**. F_decay_ kinetics for NX^Rho^C +/- 200/500 μM lidocaine at -30 mV (Welch’s t-test).

## Discussion

In this study we adapt our previously developed tPTS-based approach to introduce a rhodamine-based fluorescent probe for VCF in hNa_V_1.5 DIII-DIV linker. This semi-synthetic strategy allows us to robustly express a homogenous population of hNa_V_1.5 labelled with a bright and photo-stable fluorescent probe in its ICD, thereby circumventing the limitations of previous methodologies based on cysteine-reactive probes or non-canonical fluorescent amino acid incorporation. This allows us to record robust fluorescence changes that reflect conformational changes in the DIII-IV linker, while simultaneously monitoring the state of the hNa_V_1.5 gate through electrophysiological recordings. Importantly, this can be done in living cells and with millisecond time resolution. We reveal that upon depolarization, the fluorescent signal initially increases and remains stable during the entire voltage pulse, consistent with the displacement toward a more hydrophobic environment of the DIII-DIV linker, which is in line with both functional and structural data ^26–28,3,4,29^. Next, we show that the voltage dependence of the conformational changes observed overlaps with that of SSI and that the F_on_ kinetics match those of I_inact_. This demonstrates that we track conformational changes related to both fast inactivation and SSI and that these two transitions are likely to lead to a similar displacement of the DIII-DIV linker into a hydrophobic pocket. Interestingly, the relative kinetics of F_on_ versus I_inact_ differ from a recent rat Na_V_1.4 VCF study, where a fluorescent non-canonical amino acid incorporated into the α-helix near the IFM particle displayed a F_on_ that developed faster than I_inact_^22^. This discrepancy likely reflect distinct kinetics and voltage-dependences of Na_V_1.4 versus Na_V_1.5 and highlight subtle Na_V_ subtype and specie differences^30–32^. Overall, our methodology establishes an exquisitely sensitive sensor of conformational changes related hNa_V_1.5 inactivation that can be exploited to study the impact of various other stimuli.

To this end, we next demonstrate that iFGF12A binding to the Na_V_1.5 CTD, as evidenced by late current reduction, directly affects the kinetics of DIII-DIV linker conformational changes. This is consistent with a proposed allosteric interplay between VSDs, DIII-DIV linker and CTD during inactivation^11^. The slowing of both F_on_ and F_decay_ suggests that iFGF12A acts as a kinetic brake to modulate inactivation and that this stabilizing influence may contribute to the observed late current reduction. We further show that lidocaine-mediated channel use-dependent block does not prevent the inactivation-related DIII-DIV linker conformational change to happen. Yet 500 μM lidocaine accelerates F_on_ and slows F_decay_, suggesting that pore-blocking drugs allosterically modulate DIII-DIV linker kinetics and thereby affect inactivated state stability. The slower F_decay_ is consistent with the notion of lidocaine delaying inactivation recovery^25^. Overall, these results highlight how PPIs and pore-targeting drugs can allosterically impact the dynamic properties of the DIII-DIV linker conformational changes and thus the stability of the inactivated state. It also establishes NX^Rho^C as a valuable tool to study Na_V_1.5 intracellular conformational dynamics in response to other stimuli.

In conclusion, our tPTS-based semi-synthetic approach provides a technical advance to introduce bright and photostable fluorescent probes into membrane proteins, allowing to monitor ion channel ICD conformational changes *via* VCF during e.g. long voltage protocols. It enables incorporation of virtually any modification into intracellular domains of membrane proteins^18,20,33^, and thus complements studies using extracellular fluorescent labels, charge immobilization experiments^30– 32,34,35^, or fluorescent non-canonical amino acid incorporation to track conformational transitions in ICDs or transmembrane domains^22,36,37^. We anticipate that our strategy, alongside recent advances in tPTS-based semi-synthetic approaches^38–42^, paves the way for similar studies on other membrane proteins.

## Methods

Human Na_V_1.5, iFGF12A and split intein-fused N and C constructs were purchased as expression-ready plasmids and injected as RNA into *Xenopus laevis* oocytes for TEVC and VCF recordings. Peptide X was synthesized by Fmoc-based solid-phase peptide synthesis. Extended Methods in SI.

### Molecular biology

Human Na_V_1.5 (canonical sequence, Uniprot reference Q14524, codon-optimized), intein constructs design corresponding to N- and C-parts of hNa_V_1.5 have been previously described^18,20^. Human iFGF12A (canonical sequence, codon-optimized, Uniprot reference P61328-1) was bought as expression-ready plasmids in pUNIV from Twist Biosciences. RNA for oocyte injection was prepared by linearization (NotI restriction enzyme) and a T7 transcription kit (mMESSAGE mMACHINE™ T7 Transcription Kit, ThermoFischer Scientific).

### Expression in *Xenopus laevis* oocytes

Oocytes lobes from *Xenopus laevis* frogs are surgically removed from the animal (anaesthetized in 0.3% tricaine, under license 2014-15-0201-00031, approved by the Danish Veterinary and Food Administration). The ovarian lobes are digested with Type I collagenase (0.5 mg/mL) to remove follicular layers in OR2 buffer (in mM: 82,5 NaCl, 2,5 KCl, 1 MgCl_2_, 5 HEPES, pH 7.4). Stage V-VI oocytes are injected (nanoliter 2010 micromanipulator, WPI) with 30 nL of a mixed solution containing cRNA encoding the N- and C-intein constructs (750 ng/nL). After 24 hours of incubation in MBS (in mM: 88 NaCl, 1 KCl, 0.4 CaCl_2_, 0.33 Ca(NO_3_)_2_, 0.8 MgSO_4_, 5 TRIS-HCl, 2.4 NaHCO_3_, pH 7.4) or ND96 (in mM: 96 NaCl, 2 KCl, 1 MgCl_2_, 1.8 CaCl_2_, 5 HEPES, pH 7.4; supplemented with 0.275 g/L sodium pyruvate, 0.09 g/L theophylline, 0.05g/L gentamycin and 0.05 g/L tetracycline), 20 nL of fluorescent peptide is injected (750 μM dissolved in deionized water). The oocytes are recorded after 48 and 96 hours of expression (incubation at 18 degrees).

### Two-electrode voltage-clamp (TEVC) recordings and voltage-clamp fluorometry (VCF) recordings

Voltage-clamped sodium currents are recorded with a two-electrode voltage-clamp amplifier (OC-725C, Warner Instruments) associated with an Axon Digidata 1550 (Molecular Devices) and pClamp10 software (Molecular Devices). Oocytes are impaled with glass microelectrodes with resistances between 0,1 and 0,8 MΩ filled with 3 M KCl solution. Oocytes are clamped at -100 mV unless stated otherwise. For VCF, oocytes are placed in a custom-made chamber with their animal pole towards the water-immersed objective (IX73 with LUMPlanFL N 40x, Olympus) of an inverted microscope. A halogen lamp (Olympus TH4-200) is used for excitation with a filter set containing ET500/20x, T515lp and ET535/30m (Chroma). The emission is detected via an avalanche photodiode (APD, Photomax200, Dagan Corporation). Baseline fluorescence is subtracted via the APD. During VCF recordings, between 10 and 50 repeats of the same protocols are collected for the same signal and are averaged automatically in pClamp10.7. ND96, ND30 and ND10 are used as recording buffers: (ND96 in mM: 96 NaCl, 2 KCl, 1 MgCl_2_, 1.8 CaCl_2_, 5 HEPES, pH 7.4; ND30 in mM: 30 NaCl, 66 NMDG, 2 KCl, 1 MgCl_2_, 1.8 CaCl_2_, 5 HEPES, pH 7.4; ND10 in mM: 10 NaCl, 86 NMDG, 2 KCl, 1 MgCl_2_, 1.8 CaCl_2_, 5 HEPES, pH 7.4). Lidocaine (Sigma-Aldrich) perfusion is performed in the same buffers used as recording solutions. Drug application is performed through an automated perfusion system (Valvebank II, Automate Scientific).

### Protocols & data analysis

For initial oocyte screening to detect current and fluorescence and for the calculation of kinetic parameters, the protocol consists of a holding potential at -100 mV, followed by three 20 ms voltage steps in 5 mV increments from -30 mV to -20 mV. To determine the voltage-dependence of activation, oocytes are held at -100 mV, and sodium currents are measured at voltage steps ranging from -80 mV to +40 mV (in 5 mV increments). To determine voltage-dependent steady-state inactivation (SSI) curves, a 500 ms pre-pulse from -100 mV to -20 mV (in 5 mV increments) is followed by a 20 ms voltage step to -20 mV. Note that for activation and SSI protocols, 10 to 20 repeats are performed and averaged. Data for current and fluorescence are simultaneously recorded. Late current is calculated during SSI protocol pulse at -30 mV and represents the mean current at 400 ms after the peak current expressed as a percentage of the maximal current. For recovery from inactivation, two protocols are used. For protocol 1, the holding voltage is clamped to -100 mV for 100 ms, followed by a primary pulse to -20 mV for 50 ms and return to the holding voltage, and then a secondary pulse to -20 mV is performed after 1 ms, this protocol is repeated 30 times with the pulse delay of the secondary pulse increasing by 1 ms with each step, i.e. extending from 1 to 30 ms. For protocol 2, the secondary pulse delay is changed by 10 ms with each step, i.e. extending from 10 ms to 210 ms. To calculate the recovery from inactivation, the fractional currents remaining comparing the secondary pulse at –20 mV with the primary pulse at -20 mV are plotted versus time, and the curves are fitted mono-exponentially to extract tau values. The recordings used for kinetic measurements are based on an average of 50 repeats of a 50 ms voltage step to -30 mV from a holding voltage at -100 mV at a frequency of 3 Hz. The same protocol is used for lidocaine application. Tau values for the kinetic measurements are extracted by mono-exponential fitting of the subsequent current and fluorescence traces on Clampfit 10.7. Statistical tests used are Welch’s t test if not otherwise stated. Raw data analysis was conducted in Microsoft Excel and PrismX (GraphPad). In Figure 1D, the baselines of the current traces have been manually adjusted to 0; in Figure 2F, the fluorescence trace has been low pass filtered (Gaussian) at 420 Hz, and in all current traces the capacitive transient currents have been masked for clarity.

## Data availability

TEVC and VCF data will be made available on the Zenodo repository.

## Acknowledgments

This project has received financial support by Lundbeck Foundation (R383-2022-165) and by the European Union’s Horizon 2020 research and innovation program under the Marie Skłodowska-Curie grant agreement No. 101109359. We thank Drs. S.A. Heusser, H.C. Chua and H.T. Kurata for helpful comments on the manuscript.

