## Supplementary Information for "Protein semi-synthesis enables real-time optical tracking of intracellular conformational changes during sodium channel inactivation"

**Author Contributions:** L.P., G.H.N., K.S. and S.A.P. designed research; L.P., M.N., I.A., V.S-B., L.M.F., J.M.C. performed research; L.P. analyzed data; and L.P. and S.A.P. wrote the paper with input from all authors.

**Competing Interest Statement:** The authors declare no competing interest.

### Extended methods

Peptide synthesis. Peptide X was synthesized following two different methodologies described as method 1 (full-length synthesis) and method 2 (synthesis in two fragments) below, and both peptide stocks have been used in this study. All starting amino acids, coupling reagents and solvents were purchased from external suppliers (Iris Biotech, Merck, Avantor) and used without further purification.

Peptide X sequence:
VKIISRKSLGTQNVYDIGVEKDHNFLLKNGLVASNCFNQQKKKLGGQDIFMTEEQKKYYNAMKKLGCI
SGDSLISLA

**Synthesis (method 1).** The 77-mer Nav1.5 peptide X was synthesized using Fmoc-based SPPS on an automated PurePep Chorus, induction heating assisted, peptide synthesizer (Gyros Protein Technologies, Tucson, AZ, USA) with 25 mL glass reaction vessel using preloaded Fmoc-L-Ala-Tentagel R HMPA resin (loading 0.18 mmol/g, RAPP Polymere). Reagents were prepared as solutions in DMF: Fmoc-protected AA (0.15 M), HCTU (0.3 M) and DIPEA (0.6 M). Sequence elongation was achieved using the following protocol: deprotection (2 × 2 min, RT, 300 rpm shaking), DMF wash (3 × 20 s, RT, 300 rpm) coupling (3 × 10 min, 50°C, 300 rpm), followed by DMF washes (3 × 20 s, RT, 300 rpm). Amino acids were triple coupled using AA/HCTU/DIPEA (ratio 1:1:2) in 6-fold excess over the resin loading to achieve efficient peptide elongation. Amino acid in position 1492 (marked in blue) was coupled as orthogonally protected Lys (**K** - Fmoc-Lys(Alloc)-OH). To prevent aggregation during solid phase synthesis, the following building blocks were used: pseudoprolines (marked in red) and backbone protected Glycine (marked in orange) (see below). All the building blocks mentioned were coupled using AA/PyBOP/DIPEA (ratio 1:1:2) in 3-fold excess over the resin loading for 2-3h at 40°C while shaking (300 rpm).

**AS** - Fmoc-L-Ala-l-Ser[PSI(Me,Me)Pro]-OH

**IS** – Fmoc- L -Ile-L-Ser[PSI(Me,Me)Pro]-OH

**G** - Fmoc-Gly-DmbGly-OH

Upon successful chain elongation of the 77-mer Nav1.5 peptide X, Alloc-protecting group was removed using tetrakis(triphenylphosphine)palladium(0) ($Pd(PPh_3)_4$) (0.2 eq.) and phenylsilane ($PhSiH_3$) (20.0 eq.) in anhydrous DCM under agitation (15 min)[1]. After repetition of the protocol, the deprotection was finalized with extensive DCM washes. Micro-test cleavage followed by LC-MS analysis was performed to confirm complete orthogonal side-chain deprotection. Finally, the 77-mer was fluorescently labelled with Rhodamine 110. *N*,*N*′-Boc-protected 5-carboxyrhodamine (*N*-Boc protected Rhodamine 110, previously synthesized[2]) was solubilized in 2 mL of NMP and pre-activated with PyBOP and DIPEA before addition to the resin bound peptide (resin/Rhodamine 110/PyBOP/DIPEA, 1:1.5:1.5:6 eq). The reaction vessel was covered with aluminum foil to avoid photobleaching, and the reaction was agitated at room temperature overnight. When the reaction was complete, the resin was washed 3x with NMP, 3x with DCM and dried.

Peptide $X^{Rho}$

Rhodamine 110
VKIISRKSLGTQNVYDIGVEKDHNFLLKNGLVASNCFNQQKKKLGGQDIFMTEEQKKYYNAMKKLGCI
SGDSLISLA

**Cleavage and purification.** The synthesized 77-mer Nav1.5 peptide X was cleaved from the resin and globally deprotected using a mixture of 92.5 % trifluoracetic acid (TFA), 2.5 % $H_2O$, 2.5 % triisopropylsilane (TIPS), 2.5 % 2,2′-(ethylenedioxy)diethanethiol (DODT) for 2 h at RT. After cleavage the peptide was precipitated in ice-cold diethyl ether and centrifuged at 4350 × *g* for 10 min at 4 °C. The resulting precipitate was re-dissolved in 50:50:0.1 ($H_2O$:MeCN:TFA) and lyophilized. Purification of the 77-mer was performed with a preparative reverse phase high performance liquid chromatography (RP-HPLC) system (Agilent Technologies, 1200 series) equipped with a reverse phase C18 column (YCM-

Actus Triat C18) and using a linear gradient (15-45% of B in 50min) with a binary buffer system of $H_2O$:MeCN:TFA (A: 95:5:0.1; B: 5:95:0.1, flow rate 20 mL/min). The collected fractions were characterized by Agilent 6470 Quadrupole LC/MS System (Agilent Technologies). The purity of the fractions was determined at 214 nm on ultra-performance liquid chromatography (UPLC) (Waters Acquity). Fractions with purity >90% were pooled and lyophilized. Finally, resulting 77-mer TFA-salt peptide was salt-exchanged to yield 77-mer Nav1.5 peptide X as HCl-salt.

**Synthesis (method 2).** Peptide X was synthesized by Fmoc-based SPPS as two fragments which were subsequently ligated using native chemical ligation as previously described [3]. Fragment 1 corresponded to the N-terminal part of the peptide containing split intein A while Fragment 2 corresponded to the C-terminal part of the peptide containing the Nav1.5 DIII-DIV fragment and split intein B. To install a rhodamine 110 on position K1492 (**K**) in Fragment 2, this lysine was incorporated with its side chain orthogonally protected with an alloc group (Fmoc-Lys(Alloc)-OH). After successful elongation, Fragment 2 was N-terminally Boc protected by reacting with di-tert-butyl dicarbonate (5 equiv) and DIPEA (10 equiv) in NMP for 90 min. The Alloc-protecting group was removed with $Pd(PPh_3)_4$ (0.25 equiv) and morpholine (20 equiv) in anhydrous DCM (30 min). The deprotection was performed three times whereafter the resin was extensively washed with DCM and NMP. *N*-$Boc_2$ protected Rhodamine 110 (previously synthesized[2], 2 equiv) was solubilized in 1 mL NMP and pre-activated with HCTU (2 equiv) and DIPEA (4 equiv) before addition to the resin bound peptide The reaction vessel was covered with aluminum foil to avoid photobleaching, and the reaction was left agitating at room temperature for 90 min. The coupling was then repeated and the resin extensively washed with NMP and DCM, before release from the resin and global deprotection.

Fragment 1: VKIISRKSLGTQNVYDIGVEKDHNFLLKNGLVASN (C-terminal thioester)

Fragment 2: CFNQQKKKLGGQDIFMTEEQ**K**KYYNAMKKLGCISGDSLISLA

**Cleavage and Purification:** The peptide was cleaved from the resin with a cleavage mix (TFA/TIPS/Phenol/DODT/$H_2O$ (88:2.5:2.5:2:5, v/v) for 3h and then precipitated in and washed with ice-cold di-ethyl ether. Peptide fragments were purified by preparative RP-HPLC on a Waters HPLC equipped with a Waters 2489 UV/Vis detector, Waters fraction collector III and Waters XBridge BEH C18 OBD Prep Column (130 Å, 5 µm, 30 x 150 mm). Samples were run at a flowrate = 37.5 mL/min. Mobile phase: A = $H_2O$, B = $CH_3CN$ and C = 99% $H_2O$ + 1% TFA. Gradient for Fragment 1: 80% A/15% B/5% C to 60% A/35% B/5% C in 20 min; Gradient for Fragment 2: 75% A/20% B/5% C to 55% A/40% B/5% C in 25 min. Fraction collection was triggered by UV intensity (λ = 210 nm).

Ligation of Fragment 1 and Fragment 2 (1 equiv of each) was performed in native chemical ligation buffer (200 mM MPAA, 40mM TCEP in 6 M guanidine-HCl, 100 mM phosphate buffer, pH 7) and incubated at room temperature overnight, upon pH adjustment to pH 7 with NaOH 1M, as described previously[3]. The final ligated peptide was purified by semi-preparative RP-HPLC purifications on a Shimadzu LC-20AT HPLC system equipped with a Shimadzu SPD20A UV/Vis detector, a Shimadzu FRC-10A fraction collector and a Vydac 208TP C8 Column (300 Å, 10 µm, 10 x 250 mm, Column Temp = 40 °C). Samples were run at a flowrate = 6.50 mL/min. Mobile phase: A = 99.95% $H_2O$ + 0.05% TFA and B = 99.95% $CH_3CN$ + 0.05% TFA. Gradient: 80% A/20% B to 55% A/45% B in 25 min.

LC-MS analysis of crude reaction mixtures and pure products was performed on a Waters Acquity H-class UPLC equipped with Waters Acquity eLambda PhotoDiodeArray detector (λ = 210-800 nm), Waters UPLC BEH C4 column (300 Å, 1.7 µm, 2.1 x 50 mm) and LCT ESI Mass Spectrometer (Column Temp = 40 °C). Samples were run using 3 mobile phases: A = $H_2O$, B = $CH_3CN$ and C = 48.75% $H_2O$ + 48.75% $CH_3CN$ + 2.5% formic acid (FA) at a flow rate of 0.5 mL/min. Gradient: 94% A/2% B/4% C to 36% A/60% B/4% C in 6.5 min.

MS analysis of purified products was performed on a Waters XEVO-G2 XS Q-TOF mass spectrometer equipped with a Waters UPLC BEH C4 column (300 Å, 1.7 µm, 2.1 x 50 mm) (Column Temp = 60 °C). Samples were run with a 1.6-minute gradient (run time 3 min) using two mobile phases: D = 99.9% $H_2O$

+ 0.1% FA, E = 99.9% $CH_3CN$ + 0.1% FA (flow rate = 0.6 mL/min). Gradient: 94% A + 2% B + 4% C to 96% B + 4% C in 1.6 min. Data processing was performed using Waters MassLynx Mass Spectrometry Software 4.2 (deconvolution with Maxent1 function).

Finally, in both cases, peptide solutions (750 µM in deionized water) are aliquoted, kept in the dark and stored in -20 Celsius degrees and used on the day of the injection.
